# Presenilin and APP regulate synaptic kainate receptors

**DOI:** 10.1101/2022.02.03.478926

**Authors:** Gael Barthet, Ana Moreira-de-Sá, Pei Zhang, Jorge Castanheira, Adam Gorlewicz, Christophe Mulle

## Abstract

Kainate receptors (KARs) form a family of ionotropic glutamate receptors which regulate the activity of neuronal networks by both pre- and post-synaptic mechanisms. Their implication in pathologies is well documented for epilepsy. The higher prevalence of epileptic symptoms in Alzheimer disease (AD) patients questions the role of KARs in AD. Here we investigated whether the synaptic expression and function of KARs was impaired in mouse models of AD. We addressed this question by immunostaining and electrophysiology at synapses between mossy fibers and CA3 pyramidal cells, in which KARs are abundant and play a prominent physiological role. We observed a decrease of the immunostaining for GluK2 in the stratum lucidum in CA3, and of the amplitude of synaptic currents mediated by GluK2-containing KARs in an amyloid mouse model (APP/PS1) of AD. Interestingly, a similar phenotype was observed in CA3 pyramidal cells with a genetic deletion of either presenilin or APP/APLP2 as well as in organotypic cultures treated with γ-secretase inhibitors. Finally, the GluK2 protein interacts with full-length and C-terminal fragments of APP. Overall, our data suggest that APP stabilizes KARs at synapses, possibly through a trans-synaptic mechanism, and this interaction is under the control the γ-secretase proteolytic activity of presenilin.

## Introduction

Kainate receptors (KARs) form a family of ionotropic glutamate receptors, related to, but distinct from AMPA and NMDA receptors, which are composed of the tetrameric assembly of subunits (GluK1-GluK5) encoded by 5 genes (*grik1*-*grik5*) (Contractor, Mulle & Swanson, 2011; Lerma & Marques, 2013). KARs regulate the activity of neuronal networks by acting at pre- and postsynaptic levels, through either an ionotropic or a metabotropic action (Mulle & Crépel, 2021). KARs are abundantly expressed in the brain, in which heteromeric GluK2/GluK5 KARs appear to be a major isoform (Wenthold *et al*., 1994). The physiological role of KARs has been extensively studied at hippocampal mossy fiber (Mf) synapses between the dentate gyrus granule cells and CA3 pyramidal cells (Carta *et al*., 2014a). KARs act at a presynaptic level to regulate short- and long-term synaptic plasticity and spike transfer (Contractor, Swanson & Heinemann, 2001; Sachidhanandam *et al*., 2009). In addition, postsynaptic KARs comprising the GluK2 subunit also mediate slowly decaying EPSCs of small amplitude (Castillo, Malenka & Nicoll, 1997; Mulle *et al*., 1998), which appear to be important for the temporal integration of signals during high-frequency bursts of synaptic input (Pinheiro *et al*., 2013). In CA3 pyramidal cells, KARs show a highly segregated subcellular distribution, being concentrated at Mf-CA3 synapses and excluded from other synaptic compartments or extracellular domains (Castillo, Malenka & Nicoll, 1997; Fievre *et al*., 2016). This strict compartmentalization relies on multiple mechanisms, including a stringent control of the amount of GluK2 subunit in CA3 PCs, the recruitment/stabilization of KARs by N-cadherins (Fievre *et al*., 2016), and the transsynaptic complex formed by C1ql2/3 and neurexin which binds to GluK2 and GluK4 subunits (Matsuda *et al*., 2016). KARs have been implicated in several neurological and psychiatric disorders in genetic linkage studies, although much remains to be explored for most of the diseases (Lerma & Marques, 2013). A strong link however exist between KARs and the clinically-relevant chronic phase of temporal lobe epilepsy (TLE) (Crépel & Mulle, 2015). Pathophysiological mechanisms involve KARs composed of GluK2/GluK5 which are ectopically expressed at recurrent Mf synapses onto dentate gyrus cells (Epsztein *et al*., 2005); these aberrant KARs participate in the generation of recurrent seizures in models of chronic epilepsy (Peret *et al*., 2014). Although epilepsy is frequently associated with Alzheimer’s disease (AD) (Vossel *et al*., 2017), there is currently no report on how KAR function may be affected in the context of AD.

Mutations of two presenilin (PS) paralogs PS1 or PS2 are causal to familial forms of AD (FAD) (Sherrington *et al*., 1995). PS is the catalytic subunit of the intramembrane protease γ-secretase which cleaves several type 1 transmembrane proteins (De Strooper *et al*., 1998). Substrates of γ-secretase are mostly membrane-bound polypeptides derived from the shedding of the extracellular domain of transmembrane proteins usually by members of the ADAM (A Disintegrin And Metalloproteinase) family of metalloproteinases (Barthet, Georgakopoulos & Robakis, 2012). PS is particularly known to be involved in the processing of the amyloid precursor protein (APP), producing the Aβ peptides and the APP-intracellular domain (AICD). The γ-secretase complex also promotes the cleavage of a large number of cell surface proteins found at synaptic contacts, such as trans-synaptic adhesion molecules including cadherins, ephrins ligands and receptor EphB (Barthet *et al*., 2011). PS have been reported to regulate synaptic transmission at a presynaptic site (Zhang *et al*., 2009; Barthet *et al*., 2018). The prevalence of epileptic-like phenomenon in AD and the role of KAR in epilepsy made us investigate a possible regulation of KARs in the context of AD. We combined immunostaining and electrophysiology to investigate the effect of PS and APP processing on KAR function and synaptic expression. By comparing three mouse models, an amyloid mouse model expressing a FAD form of PS1 (ΔE9), and conditional KO model of PS and APP, we identified APP and its processing by PS as strong regulators of synaptic KARs.

## Results

### Synaptic KARs are selectively downregulated at Mf-CA3 synapses in APP/PS1 mice

GluK2 containing KARs are highly enriched at hippocampal Mf-CA3 synapses (Fievre *et al*., 2016; Carta *et al*., 2014a), as illustrated by the abundance of immunofluorescence signal in the stratum lucidum of wild-type but not GluK2^-/-^ mice (Figure 1A,B). Using GluK2/3 and GluK5 selective antibodies (see Methods), we performed immunolabeling of hippocampal sections to compare KAR expression between WT and APP/PS1. We detected a decrease in the immunofluorescence signal for GluK2 in the stratum lucidum of 6-month-old APP/PS1 mice as compared to age-matched control mice (Figure 1C, D). The quantification of mean pixel intensity within the stratum lucidum showed a significant decrease in GluK2 staining (Figure 1E). However, although GluK2 combines with GluK5 subunit to form native heteromeric KARs, we did not detect any significant change in the staining for the GluK5 subunit (Figure 1F,G).

**Figure 1.**
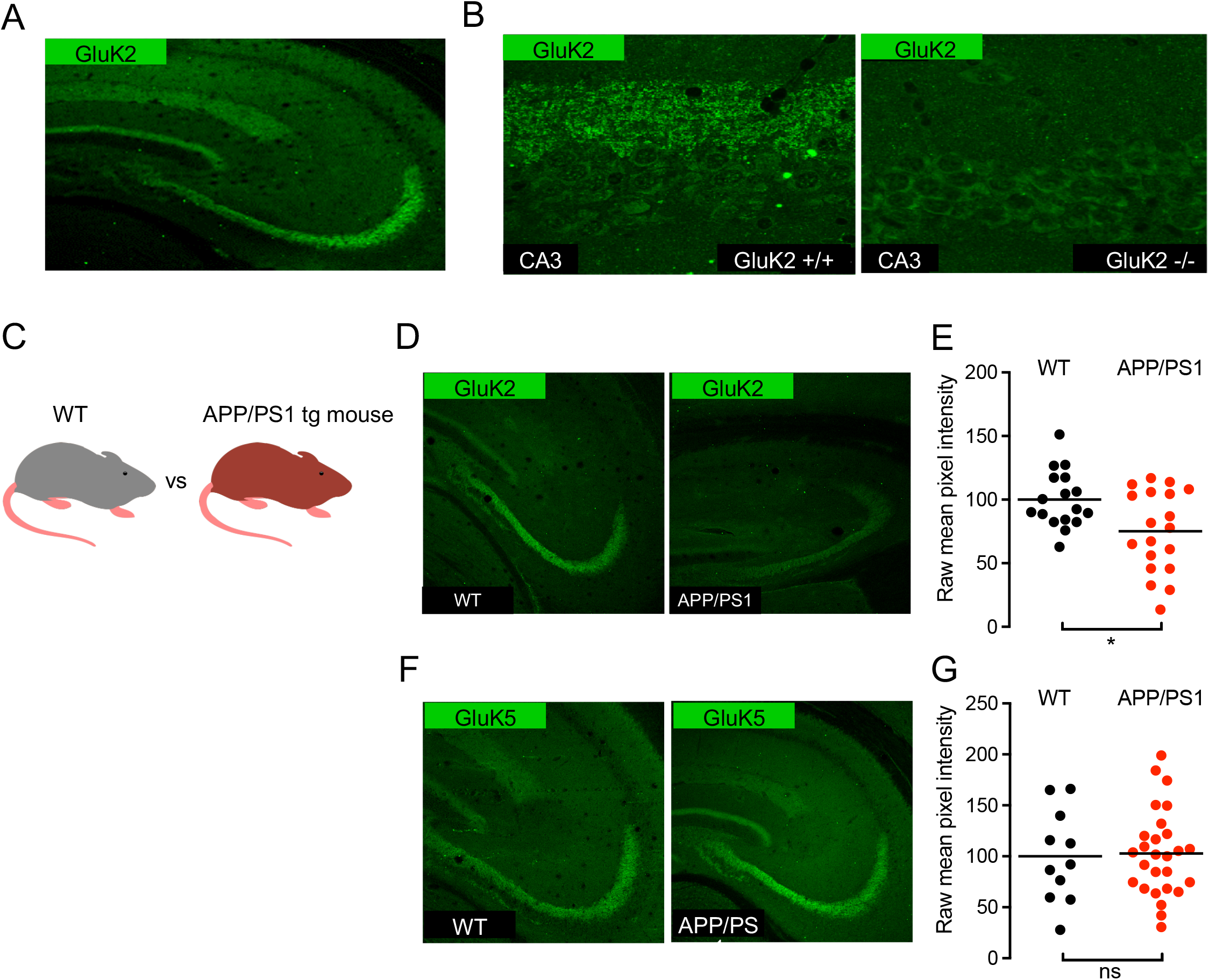
Distribution of kainate receptor subunits GluK2 and GluK5 in the hippocampus of WT and APP/PS1 mice. **A**, Immunolabelling of mouse hippocampal sections reveals strong expression of GluK2 subunit in the stratum lucidum. **B**, Zoom-in acquisitions on the CA3 area show staining adjacent to the CA3 pyramidal layer in the stratum lucidum corresponding to Mf-CA3 synapses. This staining is lost in GluK2^-/-^ mice. **C**, Cartoon representing an APP/PS1 mouse model compared to a WT littermate. **D**, GluK2 staining in the CA3 region of APP/PS1 mice is decreased in comparison to WT. **E**, Quantification of raw mean pixel intensity in the stratum lucidum region of images as in D. Data were normalized to the mean intensity of the WT condition. **F**, GluK5 staining in the CA3 region of the APP/PS1 is similar to WT. **G**, Quantification as in E shows no detectable decrease of GluK5.

We have recently shown that the basal structural and functional properties, as well as presynaptic short-term plasticity at Mf-CA3 synapses are unaltered at 6 months in the APP/PS1 mouse model of AD (Viana da Silva *et al*., 2019). At this early stage, the APP/PS1 mice however display impairment in selected forms of synaptic plasticity, namely the lipid-based DPE (depolarization-induced potentiation of excitation) (Carta *et al*., 2014b), and longterm potentiation of NMDA receptors (Viana da Silva *et al*., 2019). Using electrophysiological recordings in slices, we further tested whether synaptic KARs at Mf-CA3 synapses were impaired in 6-month-old APP/PS1 mice. We recorded from CA3 pyramidal cells in the wholecell voltage-clamp mode, in the presence of the GABA_A_ receptor blocker bicuculline (10 μM), to isolate glutamatergic inputs (Figure 2A), and the selective NMDAR antagonist D-AP5 (50 μM), to block NMDAR-mediated currents. We evoked Mf-CA3 EPSCs using minimal stimulation conditions with a glass electrode positioned close to the DG cell layer (Marchal & Mulle, 2004). As expected for Mf-CA3 synapses, the amplitude of evoked EPSCs markedly increased when the frequency of presynaptic stimulation was increased from 0.1 to 3 Hz (Figure 2B-D). Although the absolute amplitude of Mf-EPSCs was quite variable at both frequencies of stimulation, we confirmed our previous report (Viana da Silva *et al*., 2019) that frequency facilitation was not different in WT vs APP/PS1 mice (Figure 2D) (WT: 5.47 vs APP/PS1: 5.02). We then applied GYKI-53784 at 25 μM in the bath to selectively block AMPARs, and thus isolate KAR-mediated EPSCs. KAR-EPSCs represent only a small fraction of the total Mf-EPSC (less than 10 %), and can be better analyzed at a stimulation frequency of 3 Hz. We found that KAR-EPSCs were significantly smaller in 6-month-old APP/PS1 mice (Figure 2E,F), as clearly indicated by a drop in the ratio between the amplitude of KAR-EPSCs and full Mf-EPSCs (mediated by AMPAR and KARs) (5.4±0.4% to 3.0±3.9%, n=13, *t*-test, p=0.0005) between APP/PS1 and WT (Figure 2G). These results indicate that synaptic KARs are selectively downregulated at Mf-CA3 synapses in this mouse model of AD, whereas synaptic AMPARs and NMDARs are spared at least in basal stimulation conditions (see also (Viana da Silva *et al*., 2019)).

**Figure 2.**
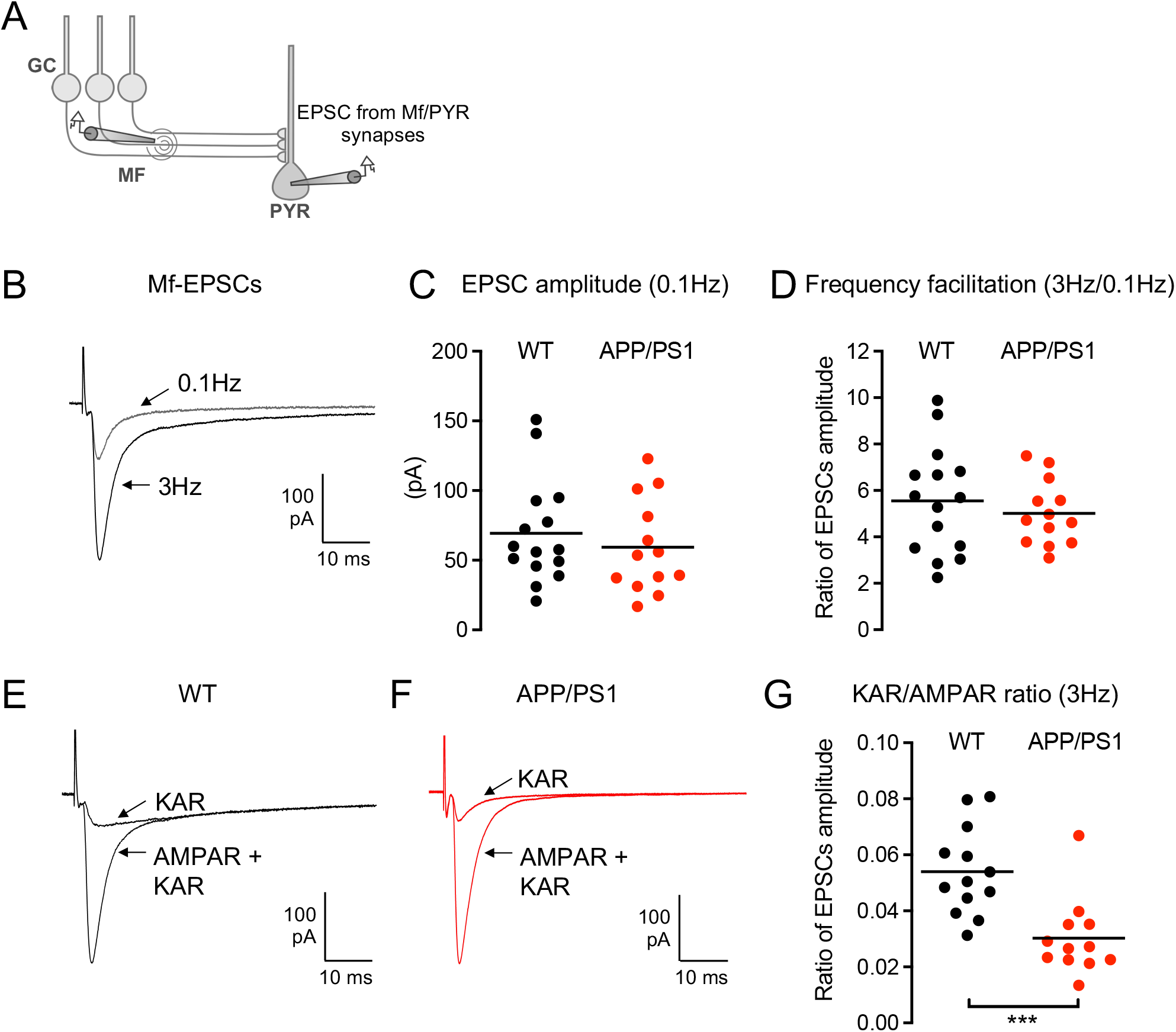
Impaired KAR-mediated synaptic transmission in APP/PS1 mice. **A**, Scheme representing the stimulation of Mf projected by granule cells (GCs) that evokes excitatory postsynaptic currents (EPSCs) recorded in CA3 pyramidal neurons (PYR). **B**, Representative traces of EPSCs recorded at a stimulation frequency of 0.1 or 3 Hz under condition in which both AMPAR and KAR currents are active. **C**, Scatter plots with averages of the EPSC amplitude recorded at 0.1 Hz demonstrate that basal synaptic transmission is not altered in APP/PS1 mice (n=15/genotype). **D**, Frequency facilitation, represented by the ratio of the amplitude of EPSCs at 3 Hz normalized to the amplitude of EPSCs at 0.1 Hz, is not altered in APP/PS1 mice. **E**, **F**, Representative traces of EPSCs recorded at 3 Hz in WT (E) or APP/PS1 (F) under conditions where both AMPAR and KAR currents are active (continuous line) or when KAR currents have been pharmacologically isolated (dashed line). **G**, Amplitude of KAR-EPSCs normalized to AMPAR-EPSCs recorded at 3 Hz. The relative amplitude of KAR-EPSCs is significantly lower in APP/PS1 mice. ***p<0.001.

### Postsynaptic deletion of presenilin decreases KAR-EPSCs at Mf-CA3 synapses

The APP/PS1 mice used in this study express the amyloid precursor protein (APP) gene with the Swedish mutations (KM670/671NL) and the presenilin 1 (PSEN1) gene with a deletion of exon 9 (Jankowsky *et al*., 2004; Garcia-Alloza *et al*., 2006). These transgenes significantly alter the γ-cleavage of APP leading to an increase in the production of aggregation-prone amyloid peptides like Aβ42 which, in turn, results in the early formation of amyloid plaques (starting at 6-months). To test whether the impaired synaptic expression of KARs resulted from an amyloid-dependent mechanism or from a more general impairment of PS-mediated transmembrane cut, we compared APP/PS1 mice to a mouse model in which PS is genetically deleted in CA3 pyramidal cells and Aβ cannot be produced. We targeted the expression of the Cre recombinase (combined with GFP) in the CA3 pyramidal cells of PS1floxed/PS2KO mice *in vivo* (Figure 3A-C). Twenty to forty days post-infection we prepared hippocampal slices and recorded Mf-CA3 EPSCs in CA3 pyramidal cells either labeled with GFP (PSdKO cells) or nonlabeled (control cells) (Figure 3D). We first evaluated potential changes in the basal synaptic properties. The amplitude of AMPAR-EPSCs recorded at 0.1 Hz was not different between the two cell types (73 pA in WT and 70 pA in PSdKO, n=11, *t*-test, p=0.8572) (Figure 3E), indicating that PS does not impact on basal content of synaptic AMPARs or on presynaptic release properties (failure rates are similar at 14%, n=11, *t*-test, p=0.9450). Unexpectedly, we found a marked decrease in AMPAR-EPSC amplitudes at 3 Hz (367 pA in WT vs 232 pA in PSdKO) (Figure 3F), hence an impairment in frequency facilitation, a characteristic form of short term synaptic plasticity at Mf-CA3 synapses (Rebola, Carta & Mulle, 2017). This impairment in short-term plasticity could either be related to the fact that the Cre recombinase is also expressed in DG cells, hence perturbing short-term plasticity at a presynaptic level (Barthet *et al*., 2018); or to a trans-synaptic consequence of deleting PS from the postsynaptic compartment. We then isolated KAR-EPSCs and normalized their amplitude to that of AMPAR-EPSCs to account for changes in glutamate release. We found that the KAR/AMPAR ratio was severely decreased, as compared to control CA3 cells (2.5%±0.4 vs 0.08%±0.3, n=11, *t*-test, p=0.0033) (Figure 3G-I).

**Figure 3.**
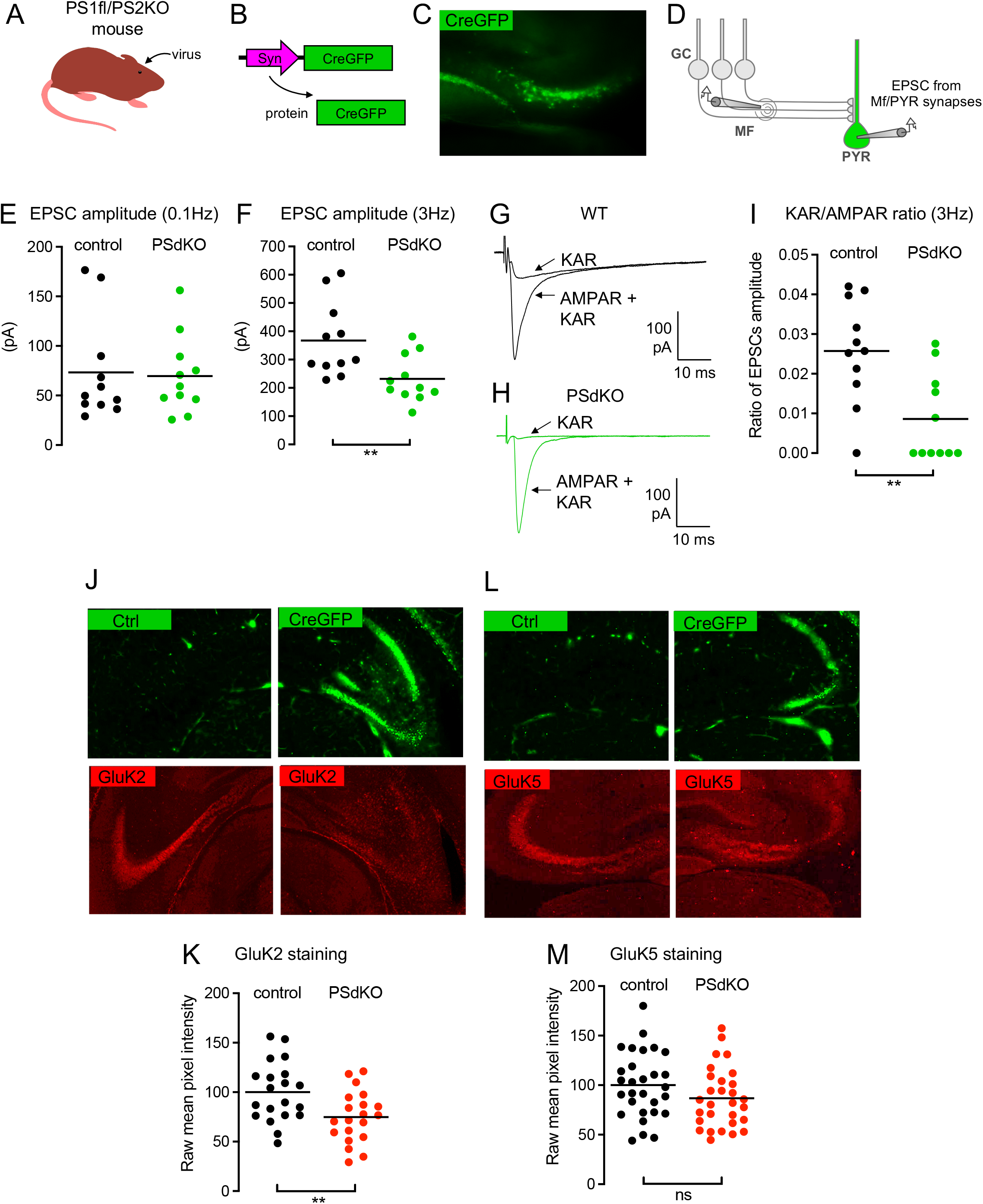
Impaired synaptic expression and function of KARs in the absence of PS. **A**, Cartoon representing the generation of PSdKO CA3 cells. Cre recombinase expressing viruses are injected in the CA3 region by stereotaxic surgery. **B**, Scheme representing the lentiviral vector to express Cre fused to GFP in CA3 pyramidal neurons. **C**, representative image of the CA3 region targeted with a virus expressing Cre-GFP. **D**, Scheme representing the stimulation of Mf that evokes EPSCs recorded in PsdKO CA3 pyramidal neurons (PYR) identified with GFP. **E**, **F**, Scatter plots with averages of the EPSC amplitude recorded at 0.1 (E) or 3 Hz (F) demonstrate that synaptic basal transmission (at 0.1 Hz) is not altered in postsynaptic PSdKO condition, whereas synaptic transmission is decreased at 3 Hz (n=15/genotype), indicating impaired frequency facilitation. **G**, **H**, Representative traces of EPSCs recorded at 3 Hz in WT (G) or in post-synaptic PSdKO (H) under conditions where both AMPAR and KAR currents are active or when KAR currents have been pharmacologically isolated. **I**, Amplitude of KAR-EPSCs normalized to AMPAR-EPSCs recorded at 3 Hz. The relative amplitude of KAR-EPSCs is significantly lower in post-synaptic PSdKO condition. **J**, GluK2 staining in the CA3 region of PS1fl/PS2 conditional KO mice targeted (right) or not (left) with a Cre-GFP-expressing virus. The staining is decreased in the post-synaptic PSdKO condition. **K**, Quantification of raw mean pixel intensity in the stratum lucidum region of images as in J. Data were normalized to the mean intensity of the WT condition. **L**, GluK5 staining in the CA3 region of the PSdKO is similar to WT. **M**, Quantification as in K shows no statistically significant decrease of GluK5. **p<0.01

We further analyzed GluK2 and GluK5 immunofluorescence labeling in the Cre-infected CA3 subregion of PS1floxed/PS2KO mice, in comparison with the non-infected contralateral CA3 (Figure 3J, L). The efficiency of infection was on the order of 60% of cells infected. We observed a consistent decrease in the staining for GluK2 in the stratum lucidum of Cre-infected CA3 as compared to contralateral CA3 (Figure 3K). Quantification of the intensity of fluorescence (see methods) showed a marked significant reduction of GluK2 labeling (but not of GluK5), even though the efficiency of infection was not maximal. Overall, our data show that presenilin controls the postsynaptic KAR content at Mf-CA3 synapses, via a mechanism which does not seem to rely on overexpression of Aβ.

### A γ-secretase inhibitor (GSI) decreases the expression of synaptic KARs

Some functions of PS have been reported to not depend on it γ-secretase-related proteolytic activity (Tu *et al*., 2006; Kallhoff-Munoz *et al*., 2008). To test whether the impairment of KAR function resulted from the loss of γ-secretase proteolysis of transmembrane proteins, we treated organotypic hippocampal slices, which preserve KAR-EPSCs at Mf-CA3 synapses (Fievre *et al*., 2016), with a γ-secretase inhibitor (GSI: L685,458 at 10 μM) for at least 4 days (from 10 DIV, after *in vitro* rewiring and synaptogenesis) (Figure 4A). We recorded from Mf-CA3 synapses and did not observe any significant change in the amplitude of the AMPAR-EPSCs at 3 Hz (Figure 4B,C). When comparing the cell-by-cell ratio of amplitudes of KAR-EPSCs vs AMPAR-EPSCs at 3 Hz, we observed a significant decrease in the KAR/AMPAR ratio in the presence of the GSI (Figure 4D)(10.1%±1.4 vs 6.6%±0.9, n=13, *t*-test, p=0.046).

**Figure 4.**
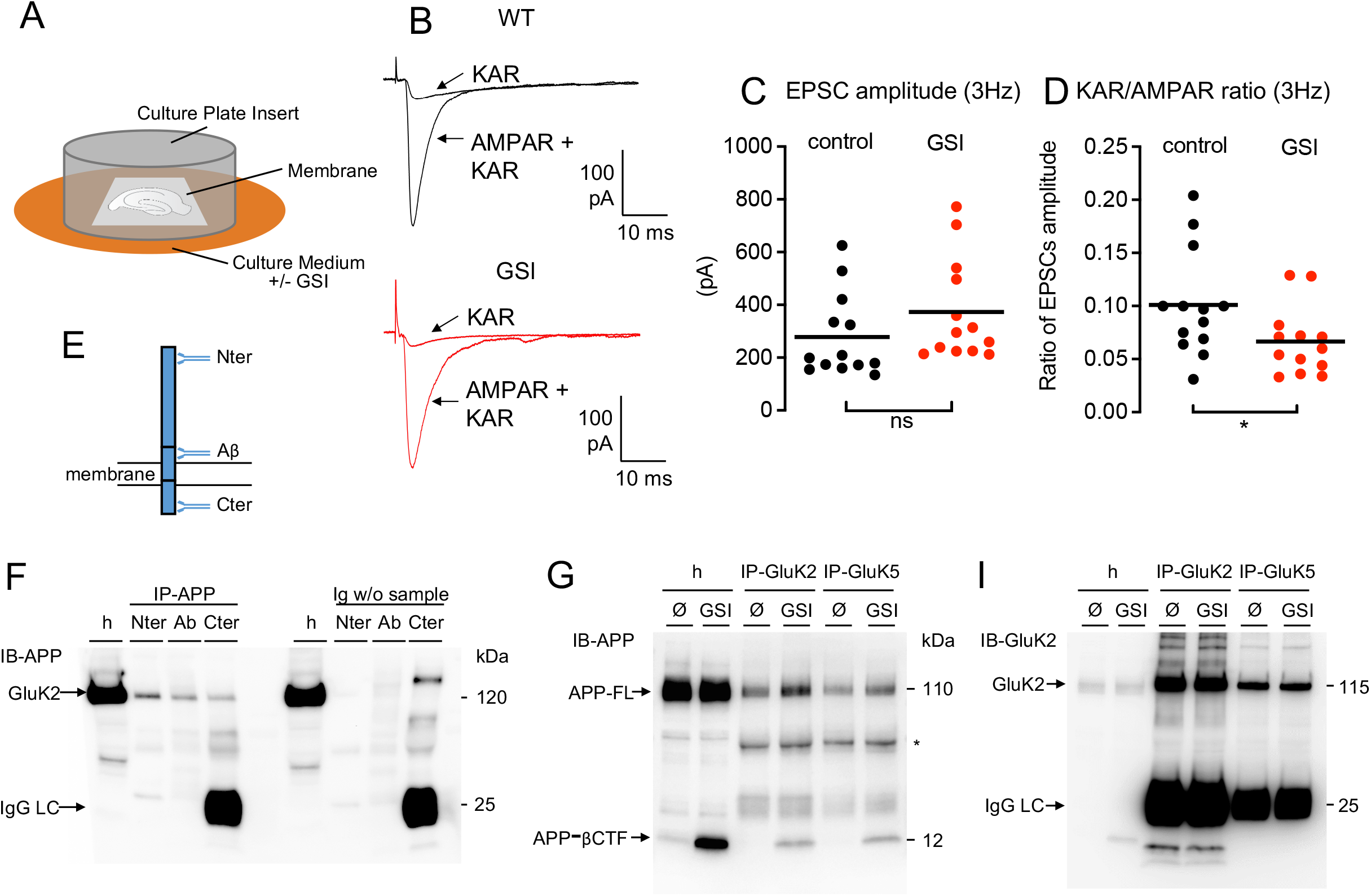
KARs are regulated by γ-secretase proteolytic function and interact with APP. **A**, Cartoon representing organotypic hippocampal slice cultures and their treatment with a γ-secretase inhibitor (GSI). **B**, Representative traces of EPSCs recorded at 3 Hz in control or in the presence of GSI, under conditions where both AMPAR and KAR currents are active or when KAR currents have been pharmacologically isolated. **C**, Scatter plots with averages of the EPSC amplitude recorded at 3 Hz demonstrate a similar AMPAR-mediated synaptic transmission between control and GSI condition (n=13, 14). **D**, Amplitude of KAR-EPSCs normalized to AMPAR-EPSCs recorded at 3 Hz. The ratio of KAR-EPSCs vs AMPAR-EPSCs is significantly lower in GSI condition. **E**, Scheme of APP full-length protein and three antibodies against three different domains (N-terminal, Aβ, C-terminal). **F**, Immunoprecipitation of APP, whatever the antibody used and the domain targeted, pulled down GluK2 subunits. The western-blot band at 120 kDa does not come from the detection of proteins from the beads or from the immuno-precipitating antibodies since the controls without sample (but with beads and immuno-precipitating antibodies) do not contain this band. **G**, Immunoprecipitation (IP) of GluK2 or GluK5, pulled down full-length APP (APP-FL) as revealed in the western-blotting of IP eluates. Treatment with γ-secretase inhibitor L685,458 causes the accumulation of APP-CTFs. In this condition, APP-CTFs co-immunoprecipitated with GluK2 or GluK5. **I**, GluK2 is highly enriched in GluK2-IP eluates. Importantly, GluK5-IP also enriches GluK2 compared to the homogenate (h), indicating strong heterodimerization of GluK2 with GluK5. *p<0.05

### Full length APP interacts with GluK2/GluK5 KARs

The regulation of KAR synaptic expression and function by the proteolytic activity of γ-secretase indicates the involvement of proteolytic substrates of PS in the regulation of KARs at synapses. We have chosen to first investigate a molecular relationship between KARs and APP. Indeed, APP is the main substrate of PS and its metabolism is particularly altered in the absence of PS (i.e. APP-CTFs accumulate in cells KO for PS or treated with GSI). Using brain homogenates, we pulled down APP by immuno-precipitation with antibodies against three different domains (N-ter, juxtamembrane, C-ter). GluK2 was co-immunoprecipitated in all three conditions, demonstrating that it likely interacts with the full-length form of APP (Figure 4E,F). In cultured hippocampal neurons, we could show that full length APP was immunoprecipitated using antibodies against both GluK2 and GluK5. We could observe that GluK2/GluK5 also interacts with the β-CTF fragment of APP, in neuronal cultures treated with GSI, in which this fragment accumulates (Figure 4G). Overall, these data indicate that GluK2/GluK5 interacts with full-length APP, possibly through a domain located in its β-CTF fragment.

### Postsynaptic deletion of APP impairs synaptic KARs

Because the level of synaptic KARs is controlled by PS and because APP, a major substrate of PS, interacts with GluK2/GluK5, we next tested whether the reduced synaptic expression of KARs upon deletion of PS was linked to APP. We targeted the expression of the Cre recombinase (combined with GFP) in the CA3 pyramidal cells of APPfl/APLP2fl mice *in vivo*. Twenty to forty days post-infection we recorded Mf-CA3 EPSCs in CA3 pyramidal cells in slices, either labeled with GFP (APP/APLP2-dKO cells) or non-labeled (control cells) (Figure 5A-C). The amplitude of AMPAR-EPSCs recorded at 0.1 Hz was not different between the two cell types (43.7±6.4 pA in WT and 55.9±5.6 pA in APP/APLP2-KO cells) (Figure 5D), indicating that APP (and or APLP2) does not impact on the basal equipment in synaptic AMPARs or presynaptic release properties (similar failure rate around 20% for both groups, n=17 for WT and n=25 for APP/APLP2-KO, p=0.9242). No change in AMPAR-EPSC amplitudes at 3 Hz (211.6±30.9 pA in WT vs 225.2±24.6 pA in APP/APLP2-KO, p=0.7304) was observed (Figure 5E), hence no change in frequency facilitation (5.28±0.8 in WT vs 4.06±0.3 in APP/APLP2-KO cells, p=0.1090). We then isolated KAR-EPSCs at 3 Hz and normalized their amplitude to that of AMPAR-EPSCs. We found that the KAR/AMPAR ratio was severely decreased, as compared to control CA3 cells (4.2%±1.0 in WT vs 1.3%±0.3 in APP/APLP2-KO, n=16-25, p=0.0022) (Figure 5F-H). This result strongly suggests that the interaction of GluK2/GluK5 KARs with APP stabilizes KARs at Mf-CA3 synapses, and that this process is under the control of the γ-secretase processing of APP by PS.

**Figure 5.**
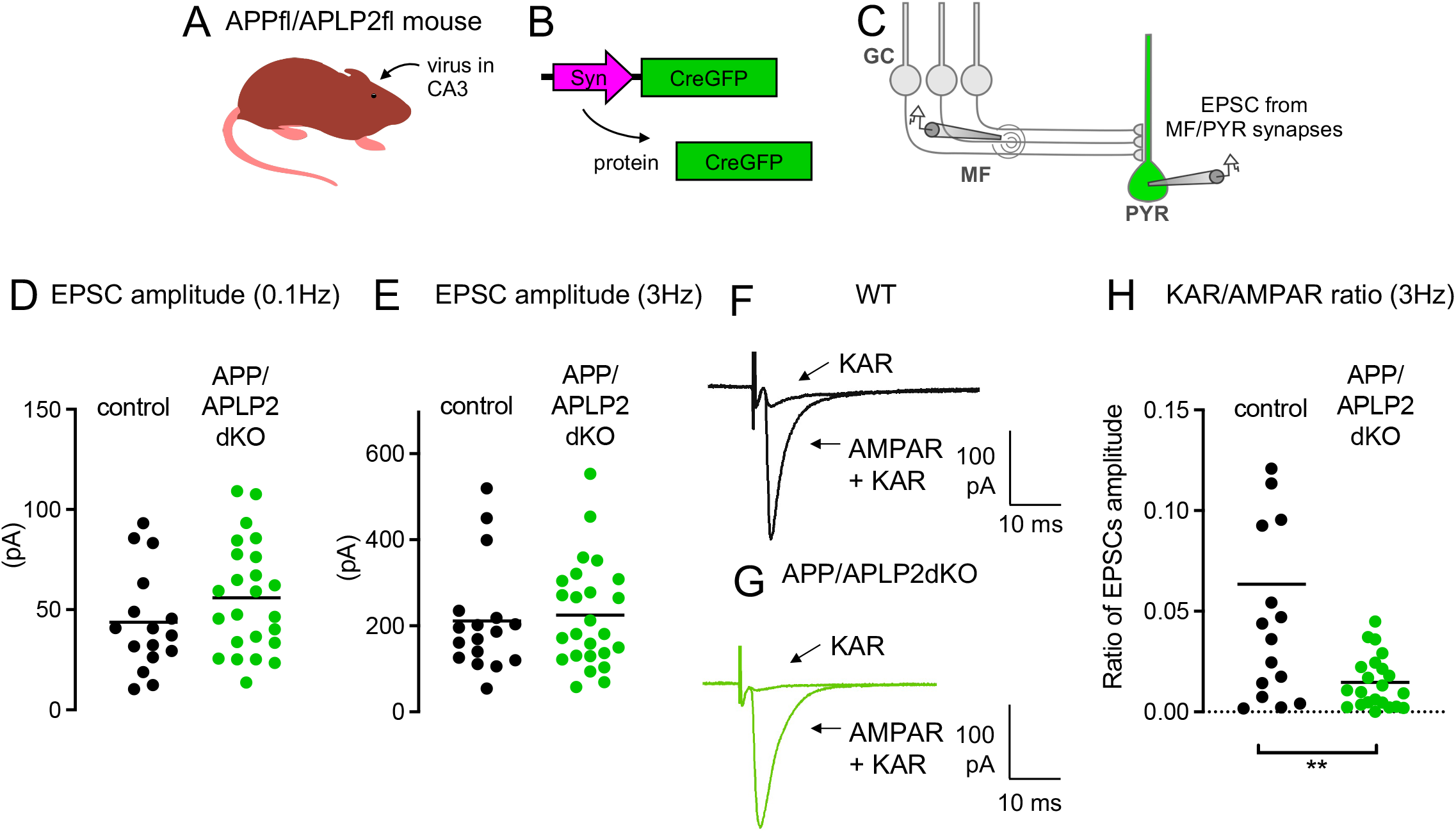
Impaired synaptic expression and function of KARs in the absence of APP. **A**, Cartoon representing the APP/APLP2 floxed mouse model. Cre recombinase expressing viruses are injected in the CA3 region by stereotaxic surgery. **B**, Scheme representing the lentiviral vector to express Cre fused to GFP in CA3 pyramidal neurons. **C**, Scheme representing the stimulation of Mf that evokes EPSCs recorded in CA3 pyramidal neurons (PYR) genetically manipulated (Cre-mediated APP/APLP2 genes deletion) and identified with GFP. **D**, **E**, Scatter plots with averages of the EPSC amplitude recorded at 0.1 Hz (E) or 3 Hz (F) demonstrate that basal synaptic transmission (at 0.1 Hz) and frequency facilitation is not altered in postsynaptic APP/APLP2 KO condition. **F**, **G**, Representative traces of EPSCs recorded at 3 Hz in WT (F) or in post-synaptic APP/APLP2 KO (G) under conditions where both AMPAR and KAR currents are active or when KAR currents have been pharmacologically isolated. **H**, Amplitude of KAR-EPSCs normalized to AMPAR-EPSCs recorded at 3 Hz. The relative amplitude of KAR-EPSCs is significantly lower in post-synaptic APP/APLP2 KO condition. **p<0.01

## Discussion

Alzheimer disease pathology involves synaptic dysfunction in its early stages (Sheng, Sabatini & Südhof, 2012). Both PS and APP, the two genes involved in the familial, early-onset form of the disease have been identified to play significant role in synaptic transmission (Müller, Deller & Korte, 2017; Shen, 2014). Understanding the regulation of synaptic receptors by PS and APP is fundamental because alteration of the physiological function of PS or APP could contribute to AD pathology (Shen, 2014). Synaptic function and plasticity have been extensively studied in mouse models in the context of AD (Marchetti & Marie, 2011; Sheng, Sabatini & Sudhof, 2012). Among these studies, the synaptic function and trafficking of AMPA and NMDA receptors was shown to be compromised in the presence of pathological concentrations of Aβ or in mouse models of AD (Sheng, Sabatini & Südhof, 2012). In contrast, there has yet been no report to our knowledge, on the link between KARs, the third family of ionotropic glutamate receptors, and AD, neither in experimental models nor in human patients.

Here we first found that immunohistological labeling GluK2 was markedly decreased in the stratum lucidum of APP/PS1 mice, at 6 months of age. The stratum lucidum corresponds to the site of synaptic contacts of mossy fiber onto the proximal dendrites of CA3 pyramidal cells. KARs comprising the GluK2 subunit are known to be expressed both at the pre- and the post-synaptic site in Mf-CA3 synapses (Mulle *et al*., 1998; Contractor, Swanson & Heinemann, 2001). At the post-synaptic site in CA3 PCs, GluK2-containing KARs are highly compartmentalized at Mf-CA3 synapses (Fievre *et al*., 2016). We then found that the amplitude of KAR-EPSCs was markedly decreased in APP/PS1 mice; in contrast, we found no change in presynaptic short-term plasticity which is known to depend on presynaptic GluK2-containing KARs (Contractor, Swanson & Heinemann, 2001). Hence the loss of GluK2 immunolabeling can be attributed to a reduced expression of KARs at the postsynaptic site. In contrast to GluK2, the immunolabeling of GluK5 did not significantly decrease in the APP/PS1 condition. This is at odds with the view that all GluK2 and GluK5 subunitsare engaged in obligatory GluK2/K5 heterodimers. Indeed, GluK5 requires heterodimerization with a GluK1-3 subunit to reach the cell surface (Ren *et al*., 2003). Our results suggests that the quantity of GluK5 subunit exceeds the pool of GluK2 leaving a significant portion of GluK5 in the trans-Golgi network unable to reach the plasma membrane. Importantly, we did not find any evidence for a change in the amplitude of AMPAR or NMDAR-mediated EPSCs in basal stimulating conditions in 6-month-old APP/PS1 mice (Viana da Silva *et al*., 2019) indicating a selective downregulation of postsynaptic KARs.

Because APP/PS1 mice overexpress a mutant form of PS (PS1Δ9), we tested whether the synaptic phenotype observed could be mimicked by deleting PS in the post-synaptic CA3 PCs. We found that the amplitude of KAR-EPSCs was similarly decreased in CA3 PCs, in parallel to a decreased immunolabeling of GluK2 in the stratum lucidum. Interestingly, a similar phenotype was observed in organotypic hippocampal cultures treated with γ-secretase inhibitors that prevent Aβ production. These experiments strongly suggested that the mechanisms underlying the control of synaptic KARs depends on PS and on its γ-secretase proteolytic activity but not on Aβ peptides.

The γ-secretase complex promotes the cleavage of a large number of cell surface proteins found at synaptic contacts, such as trans-synaptic adhesion molecules (Barthet *et al*., 2011). For example, N-Cadh is well known to undergo sequential processing, first by ADAM-10 at the juxtamembrane region leading to a membrane-tethered C-terminal fragment CTF1 (Marambaud *et al*., 2003), and then by PS at the transmembrane segment releasing the CTF2 peptide into the cytoplasm (Marambaud *et al*., 2003; Malinverno *et al*., 2010)}. We have demonstrated that N-Cadherin is involved in the synaptic stabilization of KARs (Fievre *et al*., 2016). We may thus hypothesize that the genetic or pharmacological invalidation of PS disrupts trans-synaptic N-Cadherin interaction, and impact on synaptic KARs at Mf-CA3 synapses. However,, APP, a major substrate of PS is also known to form *trans* dimers at the neuromuscular junction and at central synapses, enabling them to function as synaptic adhesion molecules (Müller, Deller & Korte, 2017). We show here that both GluK2 and GluK5, which form native heterodimers (Wenthold *et al*., 1994), form molecular complexes with both the C- and the N-terminal of APP, hence the full length protein. The genetic removal of APP (and APLP2) from post-synaptic CA3 pyramidal cells leads to a marked reduction of synaptic KAR-EPSCs. These experiments strongly suggest that APP stabilizes GluK2/GluK5 containing KARs at Mf-CA3 synapses, possibly through its role as a transsynaptic protein. This process appears to be under the control of the γ-secretase activity of PS. The disruption of γ-secretase activity may lead to a disbalance of the cleavage by-products of APP (and possibly N-Cadherin), with an increased level of the β-CTF fragment of APP, which is shown to be immunoprecipitated by GluK2 and GluK5 antibodies. KARs bound to the β-CTF fragment of APP may escape then from trans-synaptic APP, hence from synaptic sites. Whatever the mechanism our results provide strong evidence for a physiological role of APP and its metabolism by PS in regulating the recruitment and/or the stabilization of synaptic KARs. What would be the consequence of a loss of synaptic KARs at Mf-CA3 synapses in the context of AD? Because of their slow decay kinetics, EPSCs mediated by KARs are important for the integration of synaptic signals particularly during bursts of incoming inputs (Pinheiro *et al*., 2013; Sachidhanandam *et al*., 2009). This implies that in conditions of reduced PS activity, the transfer of spikes between the DG and CA3 will be impaired, in response to the typical bursting activity of the presynaptic DG cells. It is thought that DG projections to CA3 support memory through pattern separation, and may be important to the initiation of oscillations such as sharpwave ripples involved in working memory (Sasaki *et al*., 2018). Overall, the destabilization of synaptic KARs in the CA3 may participate in the difficulties in forming new memories in AD patients. Aberrant synaptic KARs expressed at aberrant recurrent fibers from DG cells (Epsztein *et al*., 2005), are critically involved in the generation of the chronic epileptic discharges in animal models of TLE (Peret *et al*., 2014). Here, we observe a reduction of synaptic KARs, which would rather limit the propagation of neuronal activity within the hippocampus. Hence, whether the PS-mediated control of synaptic KARs play any role in neuronal hyperactivity and in the epileptic propensity of AD is difficult to predict at this point. However, our study clearly indicates that KARs should be better taken into consideration whenever addressing synaptic dysfunction in the context of AD.

## Material and Methods

### Mice

The APP/PS1 mice used (Jankowsky *et al*., 2004; Garcia-Alloza *et al*., 2006) were obtained from Jackson Laboratory and used according to regulations of the University of Bordeaux/CNRS Animal Care and Use Committee. The PS1fl/PS2KO mice were produced by mating PS1floxed with PS2-KO mice lines from Jackson Laboratory (Bar Harbor, ME, USA) (ID # 007605 and 005617 respectively) (Barthet *et al*., 2018). The APP/APLP2 double floxed mice were produced by crossing APPfloxed with APLP2floxed homozygotic mice lines obtained from Ulrike Müller (Heidelberg, Germany). Throughout their life, all mice were group-housed, ranging from 4 to 10 animals per cage. Food and water were provided ad libitum. The transparent plexiglass cages (38.1 x 19.1 x 12.7 cm) were maintained on a 12 h dark/light cycle, kept in a temperature-regulated room, and protected from exterior pathogens by a filter. Experiments on APP/PS1 mice were performed in the light phase of the circadian cycle in 6 months (26-32 weeks) male and age-matched male WT littermates. Experiments on PS1fl/PS2KO and APP/APLP2 KO were performed on P40-P60 mice, three weeks after viral manipulation.

### Viral gene transfer and stereotaxic delivery

Mice (P22–P30) were anesthetized by isoflurane inhalation and injected with buprenorphine to prevent post-surgery pain. 500 nL of viral solution (1 × 10E8 particles) was injected using a micropump and syringe (Nanofil WPI) at the rate of 100 nl/min in the DG (Y: 2.2mm from lambda; X: ± 2.5 mm from sagittal suture; Z: 2.35 mm from the skull) to infect CA3 neurons. The spreading of viruses allowed the infection of a large part of the hippocampus (radius of roughly 1 mm) to produce numerous slices for IHC or electrophysiology. Experiments were performed at least 3 weeks postinjection.

### Slice preparation

Anesthesia drug (ketamine 75 mg/kg and xylazine 10 mg/kg) was diluted in saline and injected intraperitoneally to the mouse 5 min before decapitation or transcardial perfusion followed by decapitation. The head was immediately placed into a petri dish filled with ice-cold cutting solution (200mM of sucrose, 20mM glucose, 0.4 mM CaCl_2_, 8 mM MgCl_2_, 2 mM KCl, 1.3 mM NaH_2_PO_4_, 26 mM NaHCO_3_, 1.3 mM ascorbate, 0.4 mM pyruvate and 3 mM kynurenic acid, pH 7.3) oxygenated with carbogen (95 % O_2_, 5 % CO_2_). The brain was rapidly removed from the scull and parasagittal slices (350 mm) were cut with a Leica vibratome (Leica VT 1200S) in the cutting solution. The slices were then kept at 33 °C in oxygenated resting solution (110 mM NaCl, 2.5 mM KCl, 0.8 mM CaCl_2_, 8 mM MgCl_2_, 1.25 mM NaH_2_PO_4_, 26 mM NaHCO_3_, 0.4 mM ascorbate, 3 mM pyruvate and 14 mM glucose, pH 7.3) for 20 min, before being transferred to aCSF. The slices were then left at room temperature for a maximum of 6h after cutting.

### Slice culture preparation and treatment

Organotypic hippocampal slices (300 μm thickness) were prepared from P5 to P7 WT C57/Bl6J mice according to the guidelines of the University of Bordeaux/CNRS Animal Care and Use Committee. Three to 4 days after plating, the medium was replaced and then changed every 2–3 days. After 10 days in vitro, the γ-secretase inhibitor L685,458 was added at 10 μM. After a period of 4 to 6 days treatment, organotypic slices were used for electrophysiology experiments.

### Electrophysiological recordings

The recording chamber of the electrophysiology set-up was perfused with oxygenated aCSF (125mM NaCl, 2.5mM KCl, 2.3mM CaCl_2_, 1.3mM MgCl_2_, 1.25mM NaH_2_PO_4_, 26mM NaHCO_3_ and 14mM glucose, pH 7.4). CA3 pyramidal neurons in the CA3b sub-region were identified by differential interference contrast microscopy using an Olympus fixed stage upright microscope (BX51WI) equipped with a 60x magnification immersion objective at room temperature, and whole-cell patch-clamp configuration was achieved with borosilicate glass capillaries with resistance value ranging from 3 to 5 MΩ. The pipettes were filled with variant intracellular solutions depending on experiments: KMSO_3_-based solution (120 mM KCH3SO_3_, 2 mM MgCl_2_, 1 mM CaCl_2_, 20 mM KCl, 10 mM EGTA, 2 mM ATPNa_2_, 10 mM HEPES, pH 7.2) was used for current-clamp recordings, CsCl-based solution (120 mM CsCl, 2mM MgCl_2_, 2 mM CaCl_2_, 5mM EGTA, 5 mM phosphocreatine, 2 mM ATPNa_2_, 0.33 mM GTP, 10 mM HEPES, 10 mM QX314, pH 7.2) was used for IPSC recordings, and CsCH3SO_3_-based solution (100 mM CsCH3SO_3_, 3 mM MgSO_4_, 3.5 mM CaCl_2_, 20 mM EGTA, 5 mM phosphocreatine, 3 mM ATPNa2, 0.33 mM GTP, 10 mM HEPES, pH 7.2) was used for all other experiments. Cells were allowed to stabilize for 8-10 minutes after wholecell configuration was established. To monitor the access resistance during the whole recording time, a hyperpolarizing voltage step (−5 mV, 10 ms) was applied at the beginning of each trace. Series access resistance was < 20 MΩ, and when it changed by > 20 %, the recording was discarded. When held at −70 mV, neurons with a holding current > 250 pA were also rejected. Mf synaptic currents were identified according to the following criteria: 1) large paired-pulse facilitation; 2) EPSCs with a steep rising phase (~1 ms); 3) EPSCs decays free of secondary peaks that might indicate the presence of polysynaptic contamination. Liquid junction potential correction was not used for measurements of membrane potentials. Unless stated differently all baselines were established with a stimulation at 0.1 Hz and compared to the same stimulation frequency after a drug application or stimulation protocol.

Recordings were made using an EPC10.0 amplifier (HEKA Elektronik), filtered at 0.5–1kHz and analyzed using IGOR Pro and Neuromatic V2.6 software. All drugs for electrophysiological experiments were obtained from Tocris Biosciences or Sigma-Aldrich, unless otherwise stated.

### Staining of GluK2, GluK5 and GFP

Mice were initially anaesthetized with a ketamine (75 mg.kg^-1^) and xylazine (10 mg.kg^-1^) solution diluted with saline solution, being injected intraperitoneally. When the animal was deeply unconscious, it was decapitated, proceeding to remove the brain. The organ was immersed in isopentane (previously cooled with liquid nitrogen) to flash-freeze the brain. Horizontal and cryo-sectioned slices (50 μm) were obtained on Leica CM3050S Cryostat (Leica Microsystems, Germany) at −20°C. During brain slicing, slices were directly mounted on SuperFrost Microscope (VWR, USA), being then submitted to RT a few minutes to obtain some humidity on mounted slices, then preserved at −80°C until immunostaining process.

For immunohistochemistry, the slices were initially fixed with Carnoy solution (1:6 acid acetic/ethanol) for 20 min at RT, then washed one single time with PBS1X for 1 min at RT, being after permeabilized with Triton 0,1% (30 min, RT). Before the incubation of primary antibodies, a blocking solution Tween 0.05% + NBCS serum 5% was applied for 30 min at RT. Then sections were then incubated with primary antibodies. Rabbit polyclonal anti-GluK2 or anti-GluK5 antibodies (1:200) were used to label KAR subunits (Synaptic Systems: #180003 and 180103 respectively) on APP/PS1 and CA3-PS-KO hippocampal sections, and a mouse polyclonal anti-GFP antibody (1:200) was used to label Cre-GFP in order to verify virus infection in CA3-PS-KO sections. This immuno-labelling of GFP was necessary because the flash-freezing procedure damaged the 3D structure of GFP necessary to its intrinsic fluorescence. After performing three washes with PBS1X (5 min at RT), a Goat Alexa-647 Anti-Rabbit antibody (1:200) was used to detect the labelling on GluK2 or GluK5 subunits, whereas a Donkey Alexa-488 anti-Mouse antibody (1:200) was used to indicate the presence of Cre-GFP (2 hours of incubation at RT). After performing three washes with PBS1X (5 min at RT), the slices were mounted using DAPI-incorporated mounting media.

### Image Acquisition

All the images of hippocampal CA3 structures were acquired on Leica Confocal Microscope SP8-CRYO (Leica Microsystems, Germany), using a HC PL FLUOTAR 10x/0.30 dry objective, the lasers Diode405, Diode638 and OPSL488, and spectral detection adjusted for the emission of Alexa-448, Alexa-638 and DAPI, using photo-multiplicator (PMT) or hybrid detectors (HyD).

### Image Analysis

ImageJ software was used to calculate raw pixel intensity of stratum lucidum of CA3 (region of interest -RO1). The data obtained with APP/PS1 and WT littermates were normalized to the mean raw intensity in WT. In the PS mouse model, acquisitions were performed in the stratum lucidum of CA3 (region of interest where genetic deletion occurs - RO1) and in the CA1 cell layer (control region not affected by viral infection – RO2). The ratio of RO1/RO2 was calculated to each hippocampus in order to normalize the obtained data from immunostainings of PS-CA3-KO mice.

### Experimental Design and Statistical analysis

Statistical analyses were performed with Prism6 (GraphPad Software, USA). First the normality of data set was tested using the D’Agostino-Pearson omnibus normality test. If data was normally distributed, a Student’s t-test, one-way ANOVA or two-way ANOVA (analysis of variance) was performed; otherwise, nonparametric tests such as Mann–Whitney test (for unpaired data) were used. For electrophysiological data, the n values can be found in the figure legends and correspond to the number of cells analyzed. Only one recording per slice was performed. Results were presented as mean ± SEM (standard error of mean) unless stated otherwise. Statistical differences were considered significant at p < 0.05. Identification of possible outliers and consequential removal was made by the Grubb’s test.

## Bibliography

Barthet, G., Georgakopoulos, A. & Robakis, N.K. (2012) Cellular mechanisms of γ-secretase substrate selection, processing and toxicity. Progress in neurobiology. 98 (2), pp. 166–175.

Barthet, G., Jordà-Siquier, T., Rumi-Masante, J., Bernadou, F., Müller, U. & Mulle, C. (2018) Presenilin-mediated cleavage of APP regulates synaptotagmin-7 and presynaptic plasticity. Nature Communications. 9 (1), pp. 4780–14.

Barthet, G., Shioi, J., Shao, Z., Ren, Y., Georgakopoulos, A. & Robakis, N.K. (2011) Inhibitors of -secretase stabilize the complex and differentially affect processing of amyloid precursor protein and other substrates. The FASEB Journal. 25 (9), pp. 2937–2946.

Carta, M., Fievre, S., Gorlewicz, A. & Mulle, C. (2014a) Kainate receptors in the hippocampus. European Journal of Neuroscience. 39 (11), pp. 1835–1844.

Carta, M., Lanore, F., Rebola, N., Szabo, Z., Da Silva, S.V., Lourenço, J., Verraes, A., Nadler, A., Schultz, C., Blanchet, C. & Mulle, C. (2014b) Membrane Lipids Tune Synaptic Transmission by Direct Modulation of Presynaptic Potassium Channels. Neuron. 81 (4), pp. 787–799.

Castillo, P.E., Malenka, R.C. & Nicoll, R.A. (1997) Kainate receptors mediate a slow postsynaptic current in hippocampal CA3 neurons. 388 (6638), pp. 182–186.

Contractor, A., Mulle, C. & Swanson, G.T. (2011) Kainate receptors coming of age: milestones of two decades of research. Trends in Neurosciences. 34 (3), pp. 154–163.

Contractor, A., Swanson, G. & Heinemann, S.F. (2001) Kainate receptors are involved in short-and long-term plasticity at mossy fiber synapses in the hippocampus. Neuron. 29 (1), pp. 209–216.

Crépel, V. & Mulle, C. (2015) Physiopathology of kainate receptors in epilepsy. Current opinion in pharmacology. 20pp. 83–88.

De Strooper, B., Saftig, P., Craessaerts, K., Vanderstichele, H., Guhde, G., Annaert, W., Figura, Von, K. & Van Leuven, F. (1998) Deficiency of presenilin-1 inhibits the normal cleavage of amyloid precursor protein. 391 (6665), pp. 387–390.

Epsztein, J., Represa, A., Jorquera, I., Ben-Ari, Y. & Crépel, V. (2005) Recurrent mossy fibers establish aberrant kainate receptor-operated synapses on granule cells from epileptic rats. 25 (36), pp. 8229–8239.

Fievre, S., Carta, M., Chamma, I., Labrousse, V., Thoumine, O. & Mulle, C. (2016) Molecular determinants for the strictly compartmentalized expression of kainate receptors in CA3 pyramidal cells. Nature Communications. 7pp. 12738.

Garcia-Alloza, M., Robbins, E.M., Zhang-Nunes, S.X., Purcell, S.M., Betensky, R.A., Raju, S., Prada, C., Greenberg, S.M., Bacskai, B.J. & Frosch, M.P. (2006) Characterization of amyloid deposition in the APPswe/PS1dE9 mouse model of Alzheimer disease. Neurobiology of Disease. 24 (3), pp. 516–524.

Jankowsky, J.L., Fadale, D.J., Anderson, J., Xu, G.M., Gonzales, V., Jenkins, N.A., Copeland, N.G., Lee, M.K., Younkin, L.H., Wagner, S.L., Younkin, S.G. & Borchelt, D.R. (2004) Mutant presenilins specifically elevate the levels of the 42 residue beta-amyloid peptide in vivo: evidence for augmentation of a 42-specific gamma secretase. Human molecular genetics. 13 (2), pp. 159–170.

Kallhoff-Munoz, V., Hu, L., Chen, X., Pautler, R.G. & Zheng, H. (2008) Genetic dissection of gamma-secretase-dependent and -independent functions of presenilin in regulating neuronal cell cycle and cell death. The Journal of neuroscience: the official journal of the Society for Neuroscience. 28 (44), pp. 11421–11431.

Lerma, J. & Marques, J.M. (2013) Kainate Receptors in Health and Disease. 80 (2), pp. 292–311.

Malinverno, M., Carta, M., Epis, R., Marcello, E., Verpelli, C., Cattabeni, F., Sala, C., Mulle, C., Di Luca, M. & Gardoni, F. (2010) Synaptic localization and activity of ADAM10 regulate excitatory synapses through N-cadherin cleavage. [online]. 30 (48), pp. 16343–16355.

Marambaud, P., Wen, P.H., Dutt, A., Shioi, J., Takashima, A., Siman, R. & Robakis, N.K. (2003) A CBP Binding Transcriptional Repressor Produced by the PS1/ε-Cleavage of N-Cadherin Is Inhibited by PS1 FAD Mutations. Cell. 114 (5), pp. 635–645.

Marchal, C. & Mulle, C. (2004) Postnatal maturation of mossy fibre excitatory transmission in mouse CA3 pyramidal cells: a potential role for kainate receptors. The Journal of Physiology. 561 (1), pp. 27–37.

Marchetti, C. & Marie, H. (2011) Hippocampal synaptic plasticity in Alzheimer’s disease: what have we learned so far from transgenic models? Reviews in the neurosciences. 22 (4), pp. 373–402.

Matsuda, K., Budisantoso, T., Mitakidis, N., Sugaya, Y., Miura, E., Kakegawa, W., Yamasaki, M., Konno, K., Uchigashima, M., Abe, M., Watanabe, I., Kano, M., Watanabe, M., Sakimura, K., et al. (2016) Transsynaptic Modulation of Kainate Receptor Functions by C1q-like Proteins. pp. 1–17.

Mulle, C. & Crépel, V. (2021) Regulation and dysregulation of neuronal circuits by KARs. Neuropharmacology. 197pp. 108699.

Mulle, C., Sailer, A., Pérez-Otaño, I., Dickinson-Anson, H., Castillo, P.E., Bureau, I., Maron, C., Gage, F.H., Mann, J.R., Bettler, B. & Heinemann, S.F. (1998) Altered synaptic physiology and reduced susceptibility to kainate-induced seizures in GluR6-deficient mice. 392 (6676), pp. 601–605.

Müller, U.C., Deller, T. & Korte, M. (2017) Not just amyloid: physiological functions of the amyloid precursor protein family. Nature Reviews Neuroscience. 18 (5), pp. 281–298.

Peret, A., Christie, L.A., Ouedraogo, D.W., Gorlewicz, A., Epsztein, J., Mulle, C. & Crépel, V. (2014) Contribution of aberrant GluK2-containing kainate receptors to chronic seizures in temporal lobe epilepsy. CellReports. 8 (2), pp. 347–354.

Pinheiro, P.S., Lanore, F., Veran, J., Artinian, J., Blanchet, C., Crépel, V., Perrais, D. & Mulle, C. (2013) Selective block of postsynaptic kainate receptors reveals their function at hippocampal mossy fiber synapses. Cerebral Cortex. 23 (2), pp. 323–331.

Rebola, N., Carta, M. & Mulle, C. (2017) Operation and plasticity of hippocampal CA3 circuits: implications for memory encoding. Nature Reviews Neuroscience. pp. 1–13.

Ren, Z., Riley, N.J., Garcia, E.P., Sanders, J.M., Swanson, G.T. & Marshall, J. (2003) Multiple trafficking signals regulate kainate receptor KA2 subunit surface expression. The Journal of neuroscience: the official journal of the Society for Neuroscience. 23 (16), pp. 6608–6616.

Sachidhanandam, S., Blanchet, C., Jeantet, Y., Cho, Y.H. & Mulle, C. (2009) Kainate receptors act as conditional amplifiers of spike transmission at hippocampal mossy fiber synapses. 29 (15), pp. 5000–5008.

Sasaki, T., Piatti, V.C., Hwaun, E., Ahmadi, S., Lisman, J.E., Leutgeb, S. & Leutgeb, J.K. (2018) Dentate network activity is necessary for spatial working memory by supporting CA3 sharp-wave ripple generation and prospective firing of CA3 neurons. Nature Neuroscience. pp. 1–18.

Shen, J. (2014) Function and Dysfunction of Presenilin. Neurodegenerative Diseases. 13 (2-3), pp. 61–63.

Sheng, M., Sabatini, B.L. & Sudhof, T.C. (2012) Synapses and Alzheimer’s Disease. Cold Spring Harbor Perspectives in Biology. 4 (5), pp. a005777–a005777.

Sheng, M., Sabatini, B.L. & Südhof, T.C. (2012) Synapses and Alzheimer’s disease. Cold Spring Harbor Perspectives in Biology. 4 (5).

Sherrington, R., Rogaev, E.I., Liang, Y., Rogaeva, E.A., Levesque, G., Ikeda, M., Chi, H., Lin, C., Li, G., Holman, K., Tsuda, T., Mar, L., Foncin, J.F., Bruni, A.C., et al. (1995) Cloning of a gene bearing missense mutations in early-onset familial Alzheimer’s disease. 375 (6534), pp. 754–760.

Tu, H., Nelson, O., Bezprozvanny, A., Wang, Z., Lee, S.-F., Hao, Y.-H., Serneels, L., De Strooper, B., Yu, G. & Bezprozvanny, I. (2006) Presenilins form ER Ca2+ leak channels, a function disrupted by familial Alzheimer’s disease-linked mutations. Cell. 126 (5), pp. 981–993.

Viana da Silva, S., Zhang, P., Haberl, M.G., Labrousse, V., Grosjean, N., Blanchet, C., Frick, A. & Mulle, C. (2019) Hippocampal Mossy Fibers Synapses in CA3 Pyramidal Cells Are Altered at an Early Stage in a Mouse Model of Alzheimer’s Disease. The Journal of neuroscience: the official journal of the Society for Neuroscience. 39 (21), pp. 4193–4205.

Vossel, K.A., Tartaglia, M.C., Nygaard, H.B., Zeman, A.Z. & Miller, B.L. (2017) Epileptic activity in Alzheimer’s disease: causes and clinical relevance. The Lancet. Neurology. 16 (4), pp. 311–322.

Wenthold, R.J., Trumpy, V.A., Zhu, W.S. & Petralia, R.S. (1994) Biochemical and assembly properties of GluR6 and KA2, two members of the kainate receptor family, determined with subunit-specific antibodies. The Journal of biological chemistry. 269 (2), pp. 1332–1339.

Zhang, C., Wu, B., Beglopoulos, V., Wines-Samuelson, M., Zhang, D., Dragatsis, I., Südhof, T.C. & Shen, J. (2009) Presenilins are essential for regulating neurotransmitter release. 460 (7255), pp. 632–636.

